# CONSTITUTIVE STIM1L DELETION ALTERS STORE-OPERATED CALCIUM ENTRY KINETICS BUT DOES NOT PERTURB THE DEVELOPMENT AND FUNCTION OF MOUSE SKELETAL MUSCLE

**DOI:** 10.64898/2026.09.28.755229

**Authors:** Olivier Dupont, Gaetan Lopez, Elena Lilliu, Perrine Castets, Xaver Koenig, Maud Frieden, Stéphane Koenig

## Abstract

Store-operated calcium entry (SOCE) is a ubiquitous calcium (Ca^2+^) signalling mechanism that replenishes endoplasmic and sarcoplasmic reticulum Ca^2+^ stores. We previously identified STIM1L, a skeletal muscle-enriched STIM1 splice variant that enables rapid SOCE activation in human myotubes. To determine its physiological role in vivo, we generated a constitutive knockout mouse model that selectively abolishes STIM1L expression while preserving the canonical STIM1 isoform. Stim1l-KO mice developed normally and exhibited no overt abnormalities, but both sexes showed reduced voluntary wheel-running activity, suggesting neuromuscular perturbation. Skeletal muscle deprived of STIM1L did not show any histological alterations or changes in fibre-type and size. Likewise, force-frequency relationships and fatigue resistance measured ex vivo in isolated extensor digitorum longus and soleus muscles were indistinguishable from control muscles. This lack of difference may be explained by the increased expression of STIM1 in several skeletal muscles and suggests that changes in the voluntary running activity of Stim1l-KO mice may arise from STIM1L loss in non-muscular tissues, including the central nervous system. Notwithstanding, primary myotubes derived from Stim1l-KO mice displayed a marked delay in SOCE activation kinetics, whereas SOCE amplitude remained comparable to controls. This was consistent with a compensatory STIM1 upregulation maintaining overall Ca^2+^ entry capacity. Interestingly, electrically evoked Ca^2+^ transients in intact Stim1l-KO flexor digitorum brevis fibres exhibited significantly faster decay kinetics than controls. This revealed another role of STIM1L in Ca^2+^ handling in skeletal muscle that appears independent of its contribution to SOCE activation. Together, these findings establish STIM1L as a regulator of SOCE activation kinetics and indicate an additional role in myofibre Ca²⁺ clearance, while suggesting that compensatory upregulation of canonical STIM1 may preserve muscle structure and contractile function in its absence in mice.

## 1 INTRODUCTION

STIM1 is the primary calcium sensor in the endo/sarcoplasmic reticulum (ER/SR) and plays a central role in activating store-operated calcium entry (SOCE). Upon depletion of ER/SR calcium stores, STIM1 undergoes conformational changes and translocation to the plasma membrane that promote its interaction with Orai1 channels, thereby enabling calcium influx. Loss of STIM1 leads to severe immunodeficiency due to impaired lymphocyte activation, and affects other organs as well, in particular skeletal muscles. Consistently, constitutive knockout mouse models display perinatal lethality due to severe skeletal myopathy, demonstrating the essential role of STIM1 in muscle development. Moreover, a partial reduction in STIM1 expression (heterozygous mice) is sufficient to impair contractile performance under conditions requiring rapid SR calcium replenishment [1]. Skeletal muscle-specific conditional deletion of STIM1 (mSTIM1-/-) caused growth delays, reduced muscle cell proliferation, and perinatal death, underscoring STIM1’s vital role in calcium signalling for neonatal muscle growth and differentiation. During repeated tonic stimulation, calcium responses persisted in control fibres but markedly declined in mSTIM1-/-myofibres. STIM1 deletion also reduced Akt and ERK signalling, linking impaired Ca²⁺ handling to diminished activation of pathways that support muscle growth and differentiation [2]. Interestingly, tamoxifen-induced, muscle-specific STIM1 depletion in adult mice led to decreased lean mass, reduced myofibre cross-sectional area, and a shorter running distance during graded treadmill tests. This exercise intolerance was linked to faster lactate build-up and an earlier switch from fat to carbohydrate metabolism [3]. The functional importance of the STIM1-Orai1 axis in muscle physiology is further supported by models expressing a muscle-specific dominant-negative Orai1. Although excitation-contraction (EC) coupling remains largely preserved in these mice, they exhibit reduced muscle mass and pronounced fatigue, demonstrating that SOCE is dispensable for EC coupling per se but critical for sustaining repeated contractile activity [4]. Beyond skeletal muscle, STIM1 is also required for cardiac adaptation to physiological stress. Cardiac-specific STIM1 ablation prevents adaptive hypertrophy and disrupts key signalling pathways, including autophagy and AKT/mTOR signalling, in response to exercise [5]. Together, these findings establish STIM1 as a potential critical regulator of calcium-dependent signalling pathways that support both skeletal and cardiac muscle function and adaptation.

We previously identified STIM1L, a long splice variant of STIM1 predominantly expressed in skeletal muscle and upregulated during myogenesis [6]. STIM1L contains an additional exon encoding an actin-binding domain that promotes the formation of preassembled STIM1-Orai1 clusters. This structural organization enables rapid and sustained activation of SOCE, consistent with the high-frequency calcium cycling required in skeletal muscle. Thus, STIM1L represents a constitutively primed SOCE machinery optimized for tissues with high contractile demand. Loss of STIM1L should therefore impair muscle function by reducing skeletal muscle’s ability to sustain repeated calcium transients, thereby increasing fatigability. Importantly, in all STIM1-deficient mouse models, both STIM1 isoforms, the canonical form and the long isoform STIM1L, are absent. As a result, it cannot be determined whether the observed muscular effects are due to the loss of a single isoform or of both isoforms. To address this, we generated a transgenic mouse model in which STIM1L expression is specifically ablated while preserving expression of the canonical STIM1 isoform. We found that skeletal muscles in Stim1l-KO mice develop normally and show no histological or functional defects. Interestingly, Stim1l-KO mice displayed reduced voluntary running activity compared to controls. However, ex vivo force measurement suggested that this behaviour does not result from increased fatigue sensitivity. The compensatory increase in STIM1 expression in skeletal muscle may explain the lack of muscle phenotype in these mice. Consistently, SOCE amplitude was similar in Stim1l-KO and control myotubes, while it was reduced upon transient STIM1L knockdown in human muscle cells. Notably, both SOCE activation kinetics and decay of induced Ca^2+^ transients were perturbed in Stim1l-KO muscle cells, pointing to important, non-redundant roles of STIM1L in Ca^2+^ regulation.

## 2 RESULTS

### 2.1 CRISPR/Cas9-mediated generation of STIM1L-specific knockout mice

To investigate the functional role of the long isoform, STIM1L, we generated a constitutive knockout mouse using CRISPR/Cas9, specifically targeting STIM1L. To do so, we designed guide RNAs on both sides of the alternative splice site (Figure 1A, Figure S1) to delete exon 13L, specific to STIM1L, in all tissues while maintaining expression of the canonical STIM1 isoform. Under these conditions, we identified one mouse with the expected 521 bp deletion spanning the end of exon 13L and the start of intron 12 (Figure 1A). We then bred this mouse to generate the *Stim1L-KO* line. By PCR, we confirmed that Stim1l+/+ (WT) and Stim1l-/-(KO) mice carry only the wild-type and KO alleles, respectively (Figure 1B). Because STIM1L is mainly expressed in skeletal muscle, we assessed its depletion efficiency by measuring its levels in skeletal muscle. Quantitative PCR revealed that the mRNA levels of total *Stim1*transcripts (*Stim1*-pan) remain unchanged in Stim1l-KO muscle compared to WT (Figure 1C). Conversely, the transcript levels of the ubiquitous *Stim1* isoform (*Stim1*-short) were increased in Stim1l-KO mouse (Figure 1C). Western blot analysis confirmed the depletion of STIM1L in quadriceps and soleus muscles from Stim1l-KO mice (Figure 1D). Notably, there was an increase in STIM1 protein levels in quadriceps, and to a lesser extent in soleus, of Stim1l-KO mice, compared to WT (Figure 1E). This increase is consistent with the enhanced detected levels of the mRNA encoding the ubiquitous STIM1 isoform in Stim1l-KO muscles, which may directly arise from the loss of the alternative long transcript, and/or from a compensatory increase in *Stim1* transcription.

**Figure 1.**
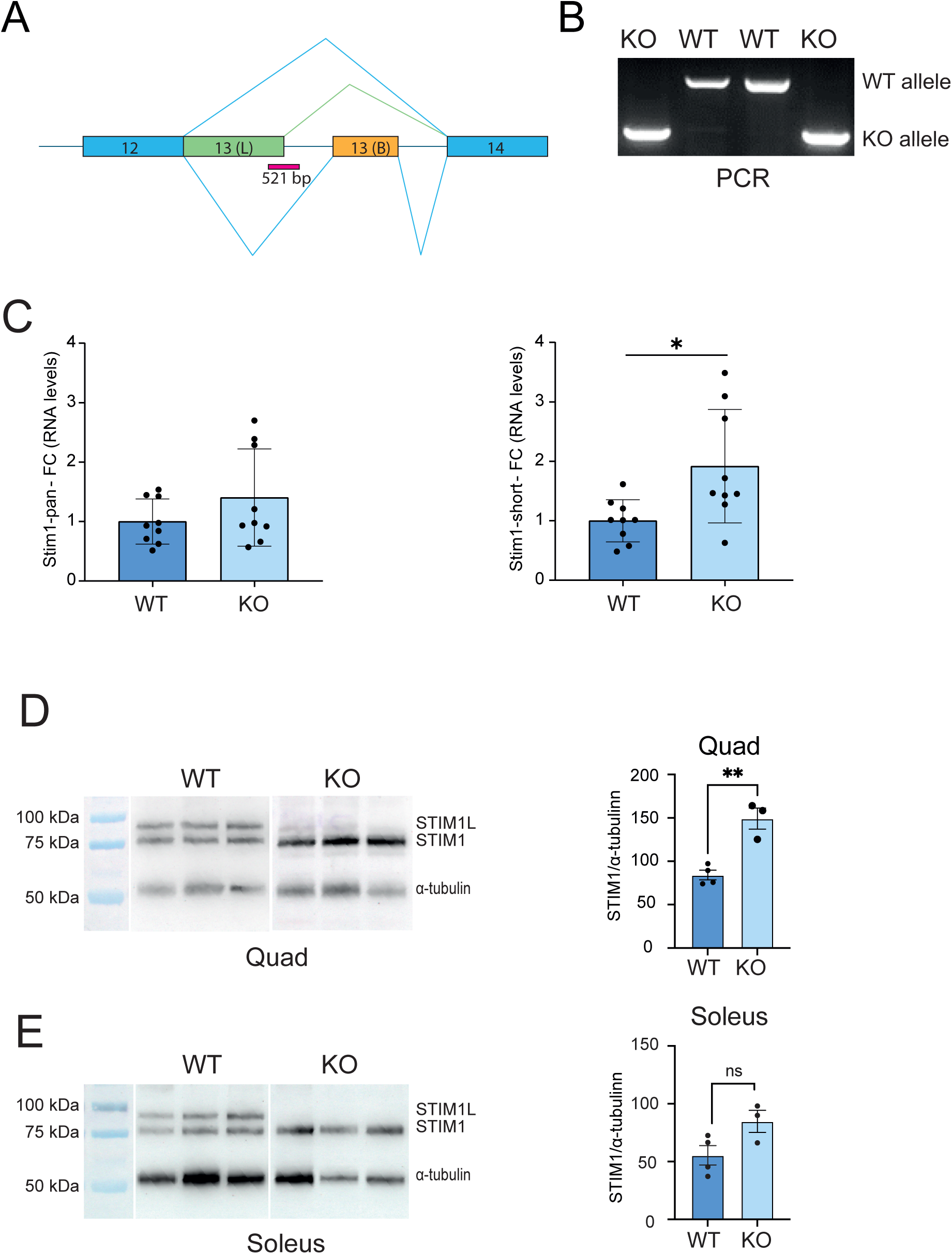
Deletion of the STIM1L-specific exon abolishes STIM1L expression and increases STIM1 expression in mouse skeletal muscle. (A) Schematic of the mouse *Stim1* gene showing alternative splicing of exon 13(L), which generates STIM1L. The deleted 521-bp region obtained in STIM1LKO mice in exon 13(L) is indicated in magenta. Exon 13B is not included in the canonical form of STIM1. (B) PCR genotyping of WT and KO mice results in products of 932 bp for the WT allele and 411 bp for the KO allele. (C) RT-qPCR analysis of total (*Stim1*-pan) and canonical (*Stim1*-short) *Stim1* transcripts in TA muscles. Expression was normalized to *Tbp* expression and expressed relative to WT. (D, E) Representative immunoblots and quantification of STIM1 protein expression in quadriceps (D) and soleus (E) muscles. α-tubulin served as a loading control. Data are mean ± [SEM/SD], with individual data points representing one mouse. Statistical significance was determined using an unpaired two-tailed Student’s *t*-test. *P* < 0.05, \**P* < 0.01; ns, not significant. KO, STIM1L knockout; WT, wild type; FC, fold change.

### 2.2 STIM1L-deficient mice develop normally but show reduced activity

Unlike Stim1-KO mice (all isoforms, [1]), Stim1l-KO mice did not display a disabling phenotype, were fertile, and showed no signs of premature mortality. At 18 weeks of age, male and female Stim1l-KO mice had the same body mass as WT mice (Figure 2A). Furthermore, EchoMRI showed no difference in lean and fat mass relative to body mass between Stim1l-KO and WT mice (Figure 2B, 2C). Overall, these data indicate that STIM1L depletion does not affect mouse development and growth.

**Figure 2.**
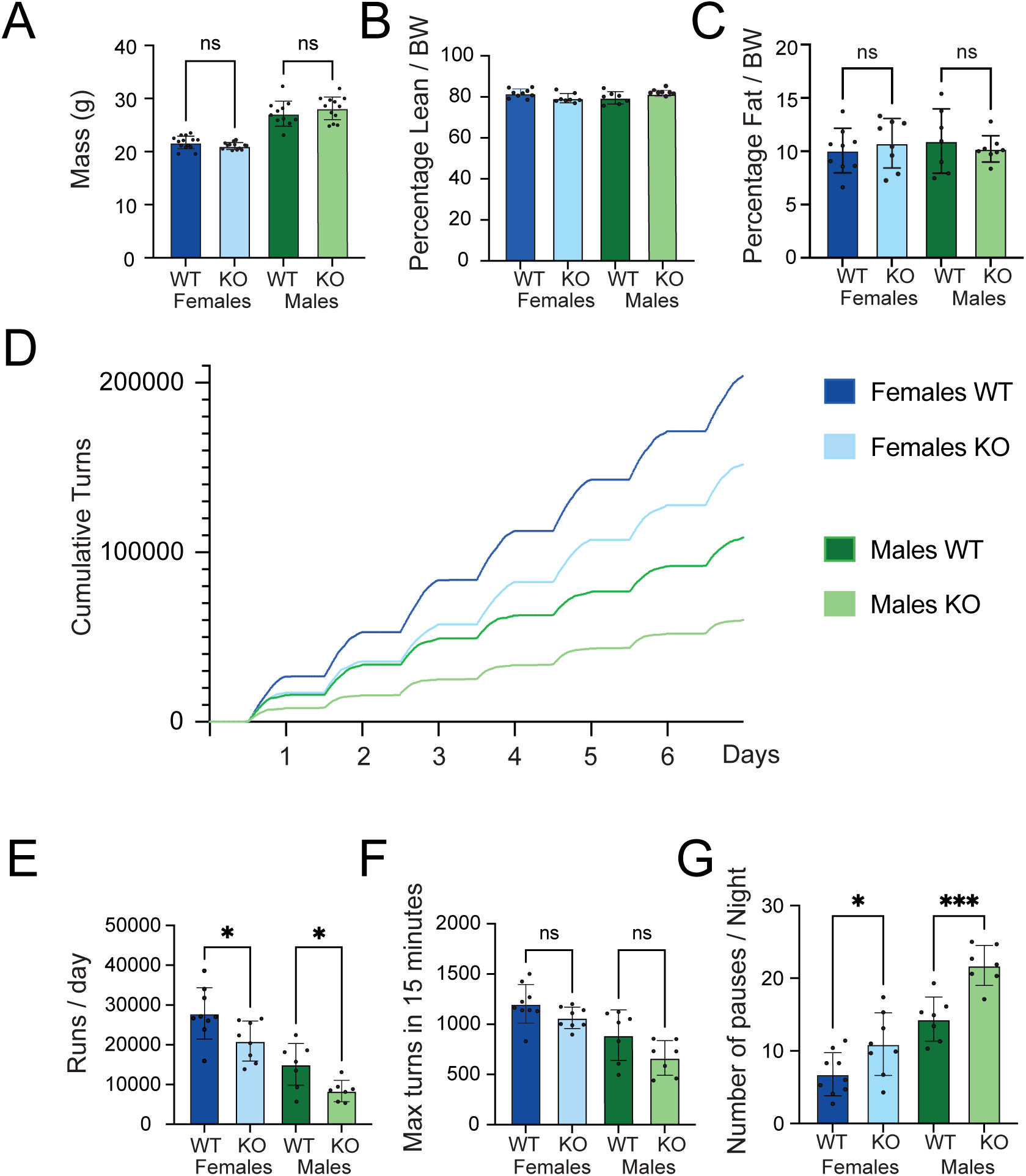
STIM1LKO mice show reduced voluntary wheel running. (A) Body mass, (B) lean mass, and (C) fat mass of female and male WT and KO mice measured by EchoMRI at 18 weeks. Lean and fat mass are expressed as percentages of body weight. (D) Mean cumulative wheel turns over 6.5 days of voluntary wheel running. Flat segments correspond to the light phase. (E) Mean daily running activity, expressed as wheel turns per day. (F) Peak running activity, defined as the highest number of wheel turns recorded in a 15-min interval. (G) Number of running pauses per night, defined as periods of ≥ 15 min without wheel activity during the dark phase. Data are mean ± SD, with individual data points representing one mouse. N = 7-9 mice per group. Statistical significance was assessed using a two-way ANOVA. P < 0.05, **P < 0.001; ns, not significant. KO, STIM1L knockout; WT, wild type; BW, body weight.

To evaluate their activity levels, we housed the mice individually in cages equipped with running wheels, enabling voluntary exercise. We found that males exhibit lower activity levels than females, regardless of genotype (Figure 2D). Interestingly, Stim1l-KO mice exhibited lower voluntary running activity, as assessed by the number of wheel turns per day, compared with WT mice in both sexes (Figure 2D). This reduction was attributable to a lower average number of wheel turns performed per day and was more pronounced in males (30% decrease) than in females (25a% decrease) (Figure 2E). Further analysis showed that the maximum number of turns performed in 15 minutes was similar between mutants and controls, indicating that Stim1l-KO mice can exert full effort comparable to WT mice (Figure 2F). However, Stim1l-KO mice took more breaks exceeding 15 minutes, which may explain the shorter distances they ran (Figure 2G). These results suggest that Stim1l-KO mice retain muscle strength but may fatigue more rapidly.

#### STIM1L-deficient mice exhibit mild muscle atrophy and a slight shift towards faster fibres

Given the reduced voluntary running activity of mutant mice and the major expression of STIM1L in skeletal muscle, we examined the muscle phenotype of Stim1l-KO mice. The absolute masses of the tibialis anterior (TA) and soleus muscles were similar in adult Stim1l-KO mice compared with controls (Figure 3A). Hematoxylin and eosin staining of TA and soleus muscle sections from Stim1l-KO mice showed no major alterations. There was in particular no sign of ongoing degeneration/regeneration, inflammation or fibrosis (Figure 3B).

**Figure 3.**
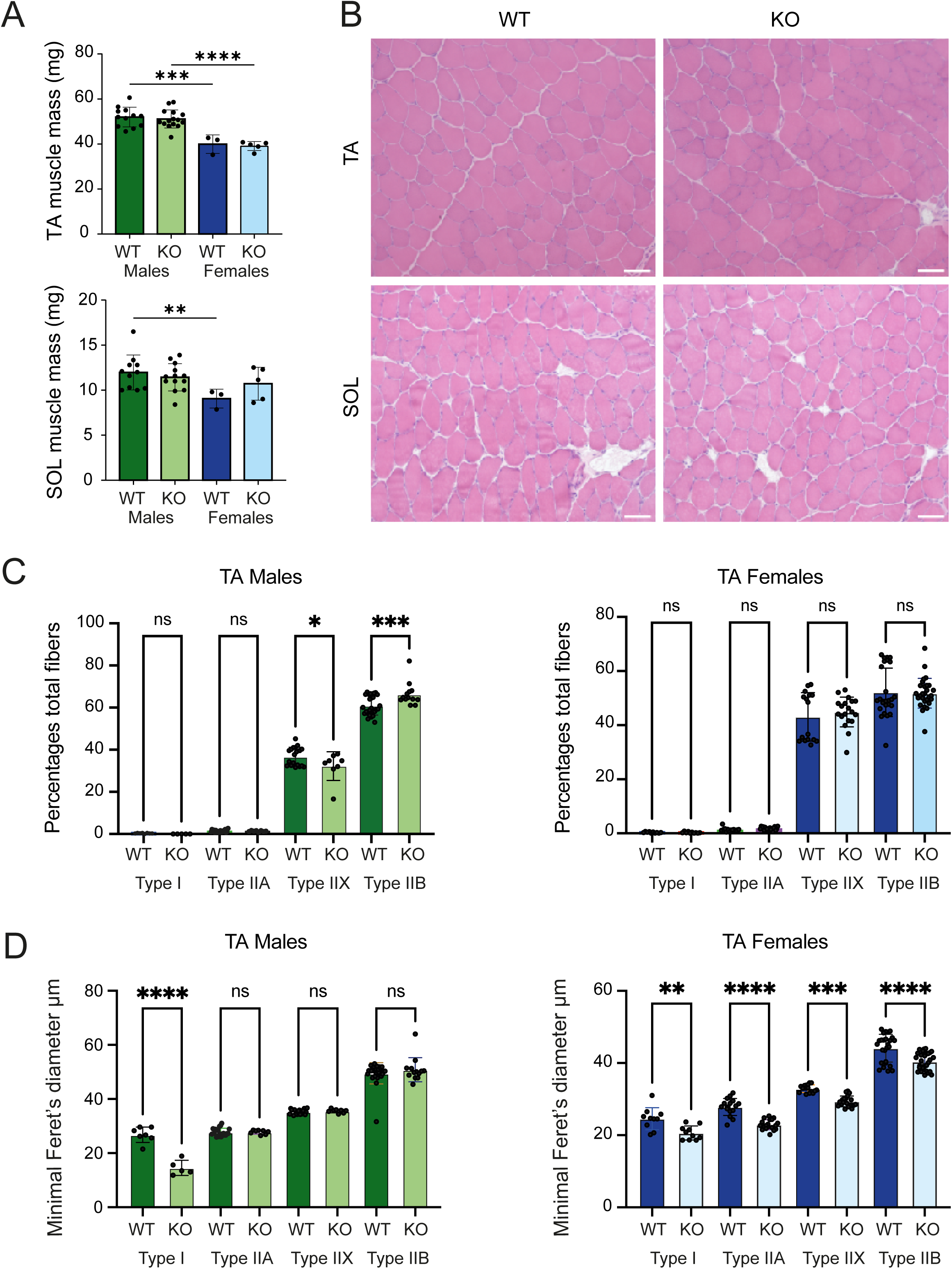
The lack of STIM1L minimally affects muscle fiber typing and does not change muscle mass. (A) Wet weight of tibialis anterior (TA, top) and soleus (SOL, bottom) from male and female WT and KO mice at 18 weeks. (B) Representative haematoxylin and eosin–stained cross-sections of TA (top) and SOL (bottom) from WT and KO mice. Scale bars, 50 µm. (C) Fibre-type distribution in TA from male (left) and female (right) mice, expressed as the percentage of total fibers that express myosin heavy chain types I, IIA, IIX, or IIB. (D) Minimum Feret diameter of each fiber type in TA from male (left) and female (right) mice. Bars represent mean ± SD. In (A), each point corresponds to one muscle. In (C) and (D), each point is from one muscle section; n = 3 mice per group. (A) was analyzed using two-way ANOVA (sex × genotype), with only significant comparisons shown. (C) and (D) were tested with an unpaired two-tailed Student’s t-test (WT vs KO). P < 0.05, *P < 0.01; ns, not significant. KO, STIM1L knockout; WT, wild type; FC, fold change; MyHC, myosin heavy chain; SOL, soleus; TA, tibialis.

To explore this further, we examined fibre-type distribution and fibre minimal Feret’s diameter in TA muscles in both males and females. As anticipated, this mainly fast muscle has very few slow-type fibres and primarily comprises type IIX and IIB fibres (Figure 3C). In females, STIM1L depletion did not alter the proportion of each fibre type compared to WT females (Figure 3C). However, Stim1l-KO males showed a slight shift from IIX to IIB fibres compared to WT (Figure 3C). Interestingly, Stim1l-KO females showed a reduction in minimal diameter across all fibre types compared with WT females (Figure 3D). In contrast, in males, STIM1L depletion affected only the size of type I fibres in TA muscle, compared with WT male mice (Figure 3D). These results indicate that STIM1L loss causes minor changes in fibre type and mild muscle atrophy in adult mice.

### 2.3 STIM1L deletion does not affect muscle force and fatigue

To directly and quantitatively evaluate how STIM1L deletion affects skeletal muscle function, we performed ex vivo tests on skeletal muscles from 18-week-old WT and Stim1l-KO mice to evaluate muscle force and fatigability. We examined the contractile properties of the extensor digitorum longus (EDL), a predominantly fast muscle, and of the soleus, which is mainly composed of slow fibres. We first stimulated the EDL and soleus muscles of both WT and Stim1l-KO mice, including males and females, with 500-ms trains across a frequency range of 1 Hz to 240 Hz and generated a force-frequency curve (Figure 4A). In these conditions, the specific forces generated (i.e., the absolute force normalized to muscle cross-sectional area, sPo) were similar for EDL muscles from WT and Stim1l-KO mice of both sexes, at all frequencies examined (Figure 4A). For soleus muscles, the plateau phase occurred at lower frequencies (60 Hz), as expected. However, there was no difference in the forces generated by Stim1l-KO and WT soleus muscles (Figure 4B). This shows that skeletal muscles lacking STIM1L, whether fast or slow, do not experience any decrease in strength compared to WT muscles.

**Figure 4.**
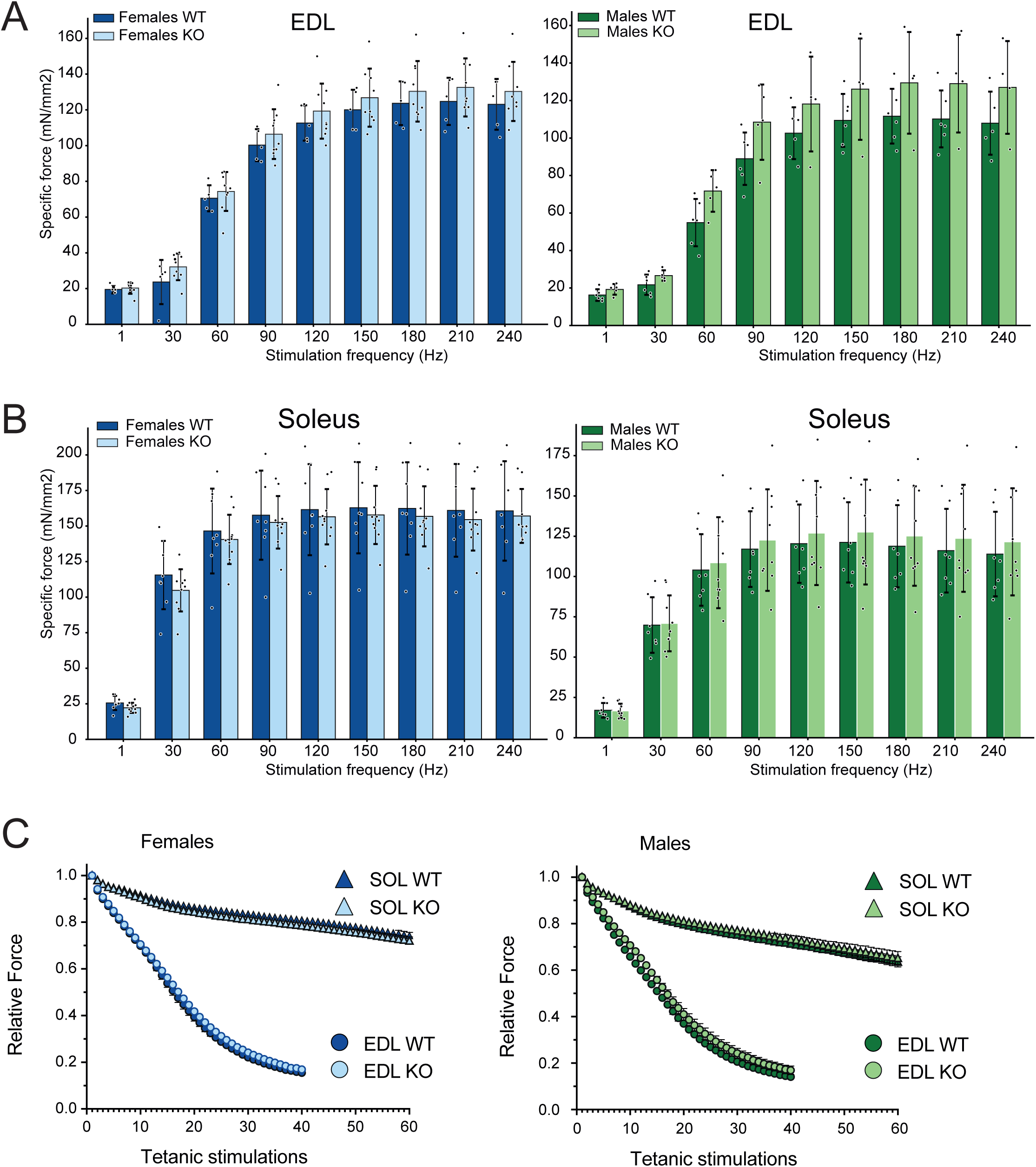
STIM1L deficiency does not impair force production or fatigue resistance in isolated fast- and slow-twitch muscles. (A,B) Force-frequency curves for isolated EDL (A) and soleus (B) muscles from female (left) and male (right) WT and KO mice. Specific force was measured ex vivo across stimulation frequencies from 1 to 240 Hz. (C) Fatigue resistance of EDL (circles) and soleus (triangles) muscles from female (left) and male (right) mice. Force is shown relative to the initial tetanic contraction. Fatigue was induced through repeated tetanic stimulations (40 for EDL and 60 for soleus). Data are presented as mean ± SD. Individual data points in (A,B) represent single muscles. EDL, extensor digitorum longus; KO, STIM1L knockout; WT, wild type.

Finally, to assess fatigue resistance in EDL and soleus muscles from Stim1l-KO mice, we applied an ex vivo fatigue protocol consisting of repeated 500-ms stimulation trains delivered every 2.5 s at 70 Hz for soleus and 130 Hz for EDL muscles. As expected, soleus muscles exhibited markedly greater fatigue resistance than EDL muscles (Figure 4C). Under these conditions, no differences in normalized force production were observed between Stim1l-KO and WT muscles throughout the fatigue protocol in either males or females (Figure 4C), indicating that fatigue resistance was preserved despite STIM1L deletion. Collectively, these results show that STIM1L loss does not impair contractile performance or alter susceptibility to fatigue during repeated muscle activity.

### 2.4 STIM1L deletion in mouse myotubes alters the kinetics, but not the amplitude of SOCE

Previous studies established that STIM1L enables rapid activation of SOCE in human skeletal muscle, which may contribute to fatigue resistance. As muscle fatigue was unchanged in Stim1l-KO mice, we next assessed the kinetics of SOCE activation in myotubes derived from myoblasts isolated from both Stim1l-KO and WT mice. SOCE was induced by rapid depletion of intracellular Ca^2+^ stores using a combination of thapsigargin and caffeine. SOCE activation was monitored using Mn^2+^ quench experiments, in which the entry of Mn^2+^ through open Orai1 channels results in a progressive quenching of Fura-2 fluorescence. Application of the thapsigargin/caffeine mixture induced a clear inflexion point, corresponding to a sharp change in the slope of the fluorescence signal (Figure 5A). We measured the time to reach the maximal quench rate as a surrogate for SOCE activation kinetics. Under these conditions, SOCE activation times were longer in Stiml1KO myotubes than in WT cells (Figure 5B). Importantly, the magnitude of this delay closely mirrors that observed in myotubes derived from human primary myoblasts transfected with either a control siRNA (siCtrl) or an siRNA targeting *STIM1L* (siSTIM1L) (Figure 5C), as previously reported [6]. These results indicate a conserved role for STIM1L in accelerating SOCE activation.

**Figure 5.**
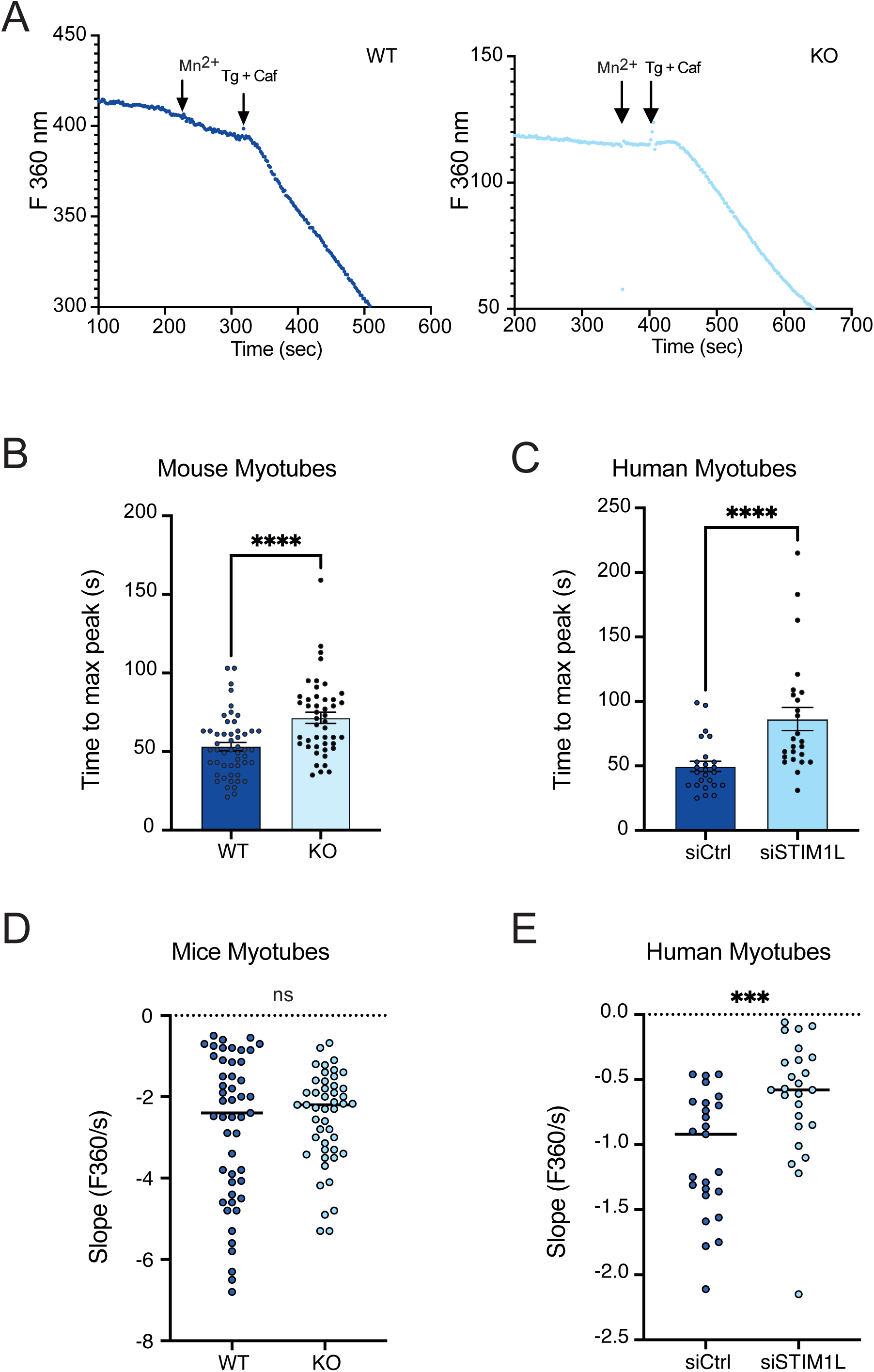
STIM1L deficiency delays store-operated Ca^2+^ entry in mouse and human myotubes. (A) Representative Mn^2+^-quench traces of Fura-2 in primary myotubes from WT (left) and KO (right) mice. Mn^2+^ (500 µM) was added first. Stores were then depleted with thapsigargin (1 µM) and caffeine (10 mM), as marked by the arrows. The rate of fluorescence quench reflects Mn^2+^ entry through store-operated channels. (B, C) Time from store depletion to the maximal quench rate, in mouse WT and KO myotubes (B) and in human myotubes transfected with control siRNA (siCtrl) or siRNA against STIM1L (siSTIM1L) (C). Mean ± SEM. (D, E) Quantification of the maximal quench rate (slope of the F360 trace) in mouse (D) and human (E) myotubes. Groups were compared by unpaired two-tailed t-test / Mann–Whitney test. ***P < .001, ****P < .0001; ns, not significant. Caf, caffeine; F360, Fura-2 fluorescence at 360 nm; KO, STIM1L knockout; siCtrl, control siRNA; Tg, thapsigargin; WT, wild type.

Given that STIM1L not only facilitates rapid SOCE activation but has also been proposed to contribute directly to SOCE activity, we next examined whether its absence affected the amplitude of Ca^2+^ entry. Because the rate of Mn²⁺-induced Fura-2 fluorescence quenching is proportional to SOCE amplitude, we quantified the maximal quenching rate in the same myotubes. Surprisingly, SOCE amplitude did not differ significantly between Stim1l-KO and WT myotubes (Figure 5D). Together with the slower SOCE activation kinetics observed in Stim1l-KO cells, these findings indicate that STIM1L deficiency delays SOCE initiation but does not affect its maximal amplitude in Stim1l-KO mouse myotubes. This observation contrasts with our in vitro findings in human myotubes, in which siRNA-mediated *STIM1L* knockdown reduces SOCE amplitude (Figure 5E). Notably, acute *STIM1L* silencing in human myotubes does not induce compensatory upregulation of the ubiquitous STIM1 isoform (Figure S2), as observed in Stim1l-KO muscle cells (Figure 1C). Hence, the reduction in SOCE amplitude in human myotubes directly reflects the loss of STIM1L-dependent Ca^2+^ entry and confirms that STIM1L contributes to total SOCE in myotubes.

We next tested whether the delayed SOCE activation and unaltered SOCE amplitude detected in STIM1L-deficient mouse myotubes also translate to mature muscle fibres. To this end, we used a skinned fibre preparation to measure SOCE as previously described [7]. In this approach, we load the transverse tubular system (t-system) of skinned EDL fibres with a Ca^2+^-sensitive fluorescent dye, enabling measurement of Ca^2+^ dynamics within this compartment. Exposure of the preparation to a release solution containing caffeine and low Mg^2+^ triggers SR Ca^2+^ release and thereby activates SOCE. This is reflected by a rapid depletion of endogenous t-system Ca^2+^ (Figure 6A). Upon re-exposure to a physiological solution containing 67 nM free Ca^2+^, the SR refills, SOCE is terminated, and t-system Ca^2+^ levels recover. Using this preparation, we detected no difference in resting t-system Ca²⁺ concentration between WT and STIM1L-deficient fibres (Figure 6A). However, t-system Ca²⁺ recovery was markedly reduced in Stim1l-KO fibres (Figure 6B). Importantly, when peak release flux (green trace, red arrow in Figure 6A) was used as a proxy for SOCE activity and normalized to the resting steady-state t-system Ca²⁺ level to account for differences in driving force, no significant difference was observed between genotypes (Figure 6C). These findings are consistent with the normal SOCE amplitude observed in intact mouse myotubes.

**Figure 6:**
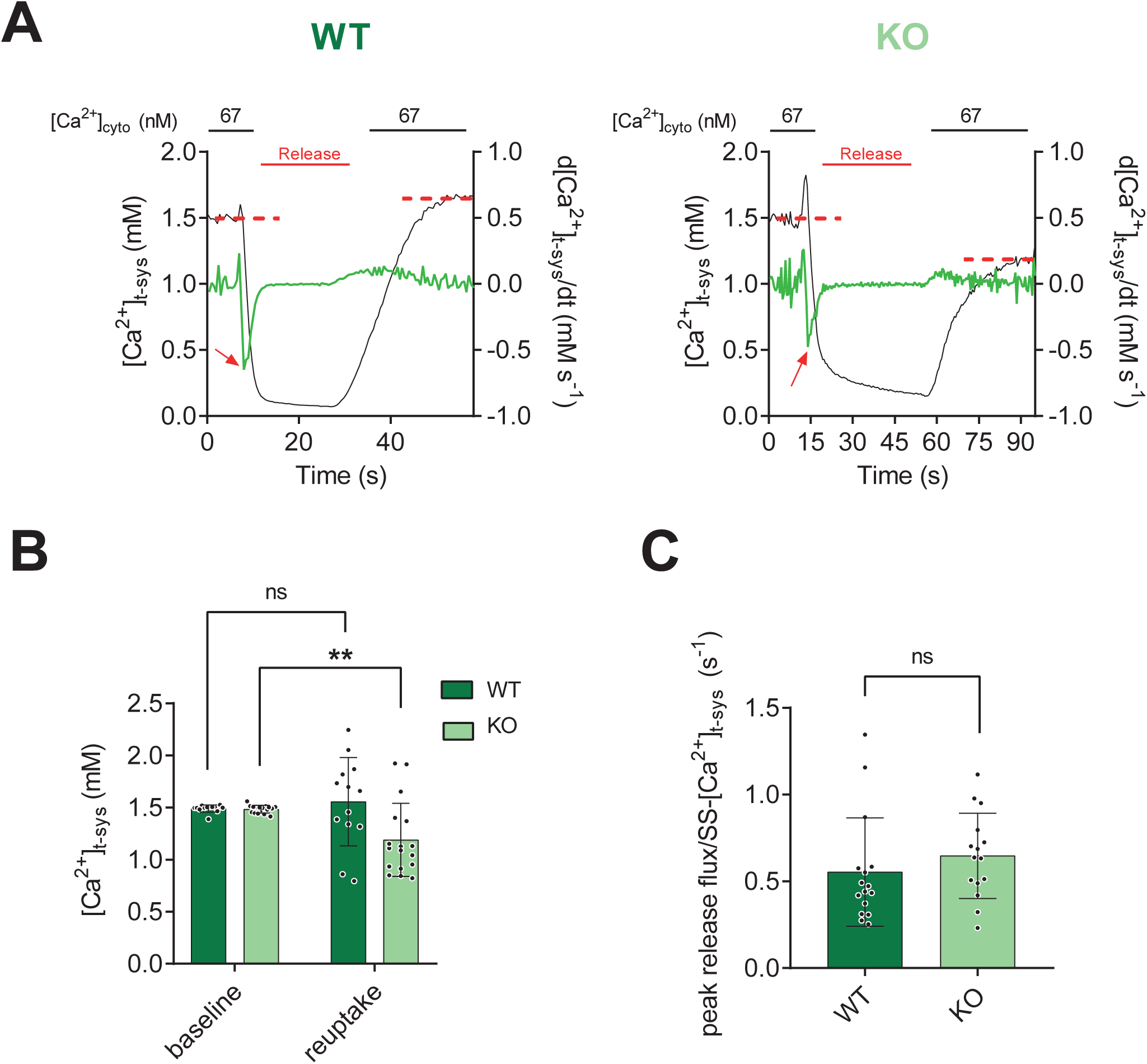
Measurement of t-system Ca^2+^ handling in skinned EDL fibers. A) Representative traces showing [Ca^2+^]_t–sys_ (black traces, left axes) as derived from the Rhod-5N fluorescence, continuously imaged during the exposure of skinned EDL muscle fibres from WT (left panel) and Stim1lKO (right panel) to a cycle of internal solutions containing 30 mM caffeine with no added Ca^2+^ and Mg^2+^(“Release”) and standard internal solution containing free [Ca^2+^] at 67 nM and free [Mg^2+^] at 1 mM. The t-system Ca^2+^ depletion (red arrows) flux was obtained by calculating the first derivative of the Ca^2+^ trace (green traces, right axes). B-C) The steady state [Ca^2+^]_t–sys_ (dashed red lines in A panels) at baseline and upon Ca^2+^ re-addition after the Release step (B) and the peaks of the [Ca^2+^]_t–sys_ depletion fluxes divided by the steady state [Ca^2+^]_t–sys_ (C) are plotted for both WT and Stim1lKO mice. Data (n= 13-16 WT and 15-17 Stim1lKO fibres) are shown as mean ± SD. Statistical significance was tested using 2-way ANOVA with Šídák’s multiple comparisons test (B) or Mann-Whitney test (C). ** = p-value < 0.01.

We next investigated whether STIM1L deficiency affects Ca²⁺ handling independently of its role in SOCE. To this end, intracellular Ca²⁺ transients were measured in intact flexor digitorum brevis (FDB) fibres during electrical field stimulation (EFS). Representative recordings from WT and Stiml1KO fibres are shown in Figure 7A. Overlaying individual Ca²⁺ transients suggested similar peak amplitudes but faster decay kinetics in Stiml1KO fibres (Figure 7B). Quantitative analysis confirmed that Ca²⁺ transient amplitude is not significantly different between WT and Stiml1KO fibres (Figure 7C), whereas decay kinetics were significantly accelerated in STIM1L-deficient fibres (Figure 7D). Calibration of resting intracellular Ca²⁺ concentrations in a subset of fibres revealed a trend towards higher resting Ca²⁺ levels in Stiml1KO fibres, although this difference did not reach statistical significance. These results confirm SOCE rescue in muscle fibres from Stim1l-KO mice, and they also suggest that STIM1L deletion may help accelerate Ca²⁺ clearance in skeletal muscle fibres.

**Figure 7:**
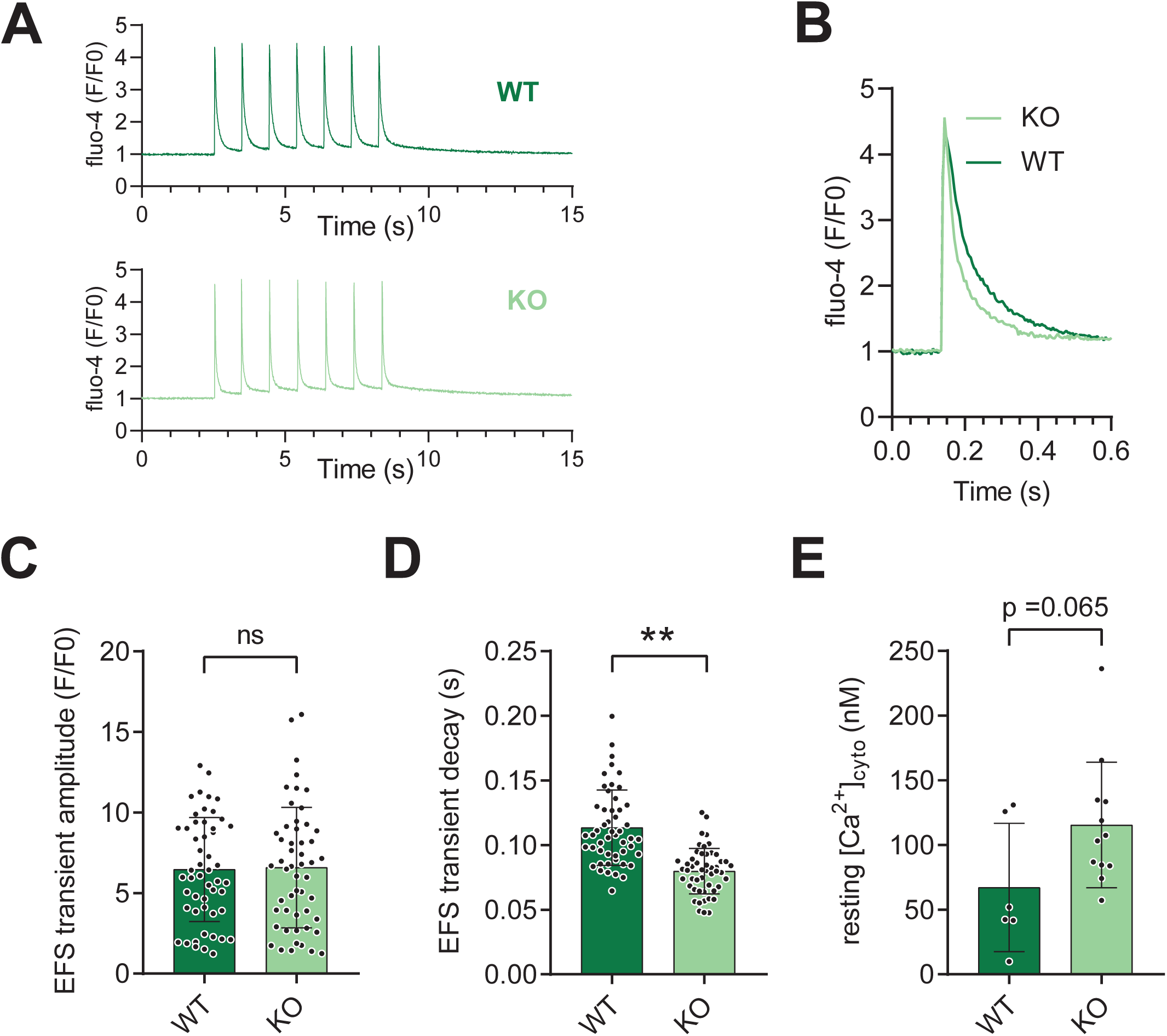
Intracellular Ca^2+^ handling in intact FDB fibers. (A) Typical cytoplasmic Ca^2+^ transients elicited by EFS of 1Hz in a WT and Stim1lKO FDB fibre. (B) Overlay of a single Ca^2+^ transient. (C, D) Statistical summary of Ca^2+^ transient amplitude and decay kinetics (time constant of exponential fit, n= 50-51 WT and 50-52 Stim1lKO fibres). (E) Statistical summary of resting cytoplasmic Ca^2+^ obtained from baseline fluorescence upon calibration (n = 6 WT and 12 Stim1lKO fibres, see Methods). Data shown as mean ± SD, statistical significance was tested using the Student’s t-test.

## 3 DISCUSSION

In this study, we generated a new transgenic mouse model lacking STIM1L (Stim1l-KO mice). We previously identified and cloned a long splice variant of STIM1, termed STIM1L, from human myotubes in vitro [6]. Unlike the ubiquitous, store-regulated isoform STIM1, STIM1L forms permanent, actin-stabilised clusters with Orai1 even when Ca^2+^ stores are full, enabling fast activation of SOCE [6], characteristic of skeletal muscle SOCE [8]. Our studies showed that this isoform is highly expressed in skeletal muscle, with lower expression in tissues such as the heart and brain [6]. Since then, other groups have identified two additional STIM1 isoforms, STIM1A and STIM1B, in other cell types, including neurons, astrocytes, and Sertoli cells [9,10]. In human myotubes, however, we found no other isoform arising from alternative splicing [6], and Western blot analysis of skeletal muscle from both WT and Stim1l-KO mice likewise revealed no additional bands (see Figure 1D and E). This confirms that the strategy used to remove STIM1L specifically did not generate a compensatory splice product.

The primary objective of generating this mouse model was to selectively abolish STIM1L expression while preserving the ubiquitously expressed STIM1 isoform, thereby enabling assessment of STIM1L’s specific contribution to skeletal muscle Ca^2+^ regulation. This model differs from all previously reported STIM1 loss-of-function mouse models, in which both STIM1 isoforms are simultaneously ablated [1–3]. Consequently, the phenotypes described in these studies cannot distinguish between functions attributable to STIM1, STIM1L, or the combined loss of both isoforms. Our *Stim1l*-specific knockout model was therefore designed to address this unresolved question. Based on our previous findings in human myotubes, where STIM1L accounts for approximately 50% of total SOCE activity, we anticipated that STIM1L-deficient muscle fibres would exhibit at least a partial reduction in SOCE, analogous to a haploinsufficiency-like phenotype [1]. Furthermore, because one of the most consistent phenotypes associated with reduced STIM1/Orai1 signalling is impaired fatigue resistance during repetitive stimulation [1,4], and as STIM1L is the isoform responsible for the rapid activation of SOCE [6], we hypothesized that exercise intolerance and muscle fatigability would constitute the predominant phenotype of Stim1l-KO mice.

Contrary to this expectation, Stim1l-KO mice displayed no overt muscular phenotype. Although both male and female knockout animals exhibited reduced voluntary running activity compared with WT littermates, they showed no evidence of muscle degeneration, structural abnormalities, or impaired resistance to fatigue. The most likely explanation for the absence of a clear muscular phenotype comes from SOCE measurements. Surprisingly, Stim1l-KO myotubes did not show the decrease in SOCE amplitude predicted by the loss of this isoform. This contrasts with our previous works in human myotubes, where acute siRNA-mediated knockdown of *STIM1L* [6,11] or shRNAmir against *STIM1L* [12] reduced SOCE by around 50%, likely because the short-term downregulation prevented compensatory upregulation of STIM1. To test this hypothesis, we repeated experiments with acute STIM1L downregulation using siRNA in primary human myotubes and observed a strong decrease in *STIM1L* expression without altering *STIM1* transcript levels. In line, SOCE decreased by around 50% under these conditions. In contrast, constitutive depletion of STIM1L in mice led to a compensatory upregulation of the ubiquitous STIM1 isoform, effectively offsetting the loss of STIM1L-dependent Ca^2+^ entry. Importantly, this compensation occurs at the transcriptional level, as Stim1l-KO muscles show increased *Stim1* transcript abundance. The underlying mechanism remains unclear. However, when examining total *Stim1* mRNA levels across all isoforms (pan-Stim1), no significant difference was found between WT and Stim1l-KO mice. These findings indicate that the overall expression of all STIM1 isoforms remains unchanged in Stim1l-KO mice. In WT mice, mRNA expression is split between STIM1 and STIM1L, whereas in Stim1l-KO mice, only the ubiquitously expressed isoform STIM1 is present, accounting for 100%.

This compensation is noteworthy given the severe phenotypes reported in models with reduced total STIM1 expression. Global STIM1 knockout mice completely lack SOCE and die perinatally from profound skeletal myopathy [1]. Even heterozygous STIM1-deficient animals display contractile defects under conditions requiring rapid SR Ca^2+^ refilling. Similarly, early muscle-specific *Stim1* ablation causes neonatal lethality associated with impaired calcineurin/NFAT, ERK1/2, and Akt signalling pathways essential for muscle growth and maturation [2]. Moreover, inducible deletion of *Stim1* in adult skeletal muscle, while circumventing developmental lethality, still results in reduced lean mass, diminished exercise capacity, altered substrate utilization, and activation of ER/SR stress and proteostatic pathways [3]. Collectively, these studies indicate that muscle dysfunction correlates closely with the extent of SOCE reduction. The absence of a marked muscle phenotype in Stim1l-KO mice is consistent with the concept that pathological consequences appear to scale primarily with the magnitude of SOCE disruption rather than the specific isoform affected [13]. Therefore, maintaining total SOCE via STIM1 compensation should maintain muscle homeostasis. Even with the full genetic absence of the actin-associated STIM1L isoform responsible for rapid SOCE activation, overall muscle function stays largely preserved as long as the net SOCE amplitude is maintained within the physiological range. Gain-of-function models further support this interpretation. Constitutively active *STIM1* mutations, which enhance SOCE, cause tubular aggregate myopathy and Stormorken syndrome in humans and induce progressive myopathy with reduced muscle force and enhanced relaxation time in knock-in mice carrying activating *Stim1* mutations such as R304W [14,15]. When considered together with the loss-of-function studies described above, our findings suggest that skeletal muscle function depends on maintaining SOCE within a relatively narrow physiological window. Deviations in either direction are deleterious, whereas preserving overall SOCE amplitude appears sufficient to sustain normal muscle mass and exercise performance.

These observations suggest that the primary physiological requirement is maintaining SOCE activity within an optimal range. However, STIM1L deficiency may also alter other aspects of Ca²⁺ homeostasis, especially in mature fibres, prompting us to investigate its role in Ca²⁺ extrusion and excitation-contraction coupling in mature skeletal muscle fibres. Quantification of Ca^2+^ extrusion at the t-tubule using skinned fibre preparations showed that this process is slowed in fibres from Stim1l-KO mice. This finding is consistent with our recent work in human myotubes, a developmentally immature in vitro model, where *STIM1* knockdown likewise impaired Ca^2+^ extrusion [12]. In that study, we identified STIM1 as the main isoform driving PMCA1-mediated Ca^2+^ extrusion in human myotubes, but *STIM1L*-specific knockdown also produced a milder effect, indicating that both isoforms contribute to this pathway, with STIM1 playing the dominant role. The slowed extrusion we now observe in Stim1l-KO skinned fibres is therefore in line with this partial, STIM1L-dependent contribution to Ca^2+^ extrusion. In apparent contradiction with this impaired extrusion, quantification of intracellular Ca^2+^ transients in FDB fibres during electrical field stimulation (EFS) revealed significantly faster, not slower, Ca^2+^ clearance kinetics in Stim1l-KO fibres compared with WT controls. In adult skeletal muscle, the cytosolic Ca^2+^ transient generated during excitation-contraction coupling is terminated predominantly by Ca^2+^ reuptake into the SR via SERCA pumps, which account for the majority of Ca^2+^ removal and largely dictate relaxation kinetics [16,17]. By contrast, PMCA-mediated extrusion across the plasma membrane contributes only marginally to net Ca^2+^ removal following a single stimulus [18,19]. The skinned-fibre and intact-fibre measurements from Stim1l-KO muscles therefore do not report on the same dominant pathway: a slowdown in the minor, PMCA-dependent extrusion component can coexist with, and be masked by, an acceleration of the major, SERCA-dependent reuptake component that determines the dynamics of the whole-fibre Ca^2+^ transient. Overall, these results show that Stim1l-KO fibres have reduced Ca^2+^ extrusion speed but increased SERCA activity. This enhanced SERCA function primarily drives the faster Ca^2+^ clearance seen during EFS. Two mechanisms could underlie this gain in SERCA function. First, STIM1L may normally exert a tonic inhibitory influence on SERCA under physiological conditions; its absence in Stim1l-KO fibres would relieve this constraint, accelerating Ca^2+^ sequestration into the SR and shortening the decay phase of the cytosolic transient. Second, and not exclusive of the first, the compensatory overexpression of STIM1 that we discuss above to explain the preserved SOCE amplitude in Stim1l-KO muscle could itself act directly on SERCA. Indeed, STIM1 has been shown to positively regulate SERCA1a in skeletal muscle [20] and SERCA2a in cardiac muscle, where it binds phospholamban and relieves its inhibitory constraint on the pump [21].

Finally, one unresolved observation remains. Reduced voluntary running activity was consistently detected in both male and female Stim1l-KO mice and represents the sole robust phenotype distinguishing mutant animals from controls. Whether this behaviour reflects a subtle muscle-intrinsic defect that escapes detection by conventional assessments of fatigue and muscle damage, or instead arises from alterations in motivation, metabolism, or central nervous system function, remains unknown. Addressing this question will require complementary behavioural, metabolic and neuromuscular investigations designed to disentangle peripheral from central determinants of exercise performance.

Our findings do not conclusively establish a specific role for STIM1L in skeletal muscle function. One possible explanation is that compensatory upregulation of canonical STIM1 obscures the contribution of STIM1L, potentially leading to an underestimation of its physiological importance in this constitutive knockout model. Defining this role will require approaches that distinguish the direct effects of STIM1L loss from compensatory adaptation and address its contribution to Ca²⁺ homeostasis beyond SOCE activation.

## 4 MATERIALS AND METHODS

### 4.1 Mice

Stim1l-KO mice were generated using CRISPR/Cas9-mediated genome editing to delete a region of the *Stim1* gene encompassing the *Stim1l*-specific splicing site (from 102080200 to 102080720, Mus musculus strain C57BL/6J chromosome 7). Two guide RNAs, gRNA1 (GGTATCTCTGCGGCGAATGC) and gRNA2 (CACGATCCTATATCACTTGT), targeting exon 13L and intron 14, respectively, were cloned into separate Cas9-encoding PX330 plasmids. These plasmids were microinjected into the pronuclei of B6D2F1 one-cell embryos. Injected embryos were subsequently transferred into pseudopregnant CD1 females. One founder mouse was obtained and crossed with C57BL/6J mice to expand the colony. Congenic C57BL/6 strains were established through successive backcrossing using IVF-assisted speed congenics (Charles River, Lyon, France). For genotyping, genomic DNA was extracted from tissue samples using the KAPA Mouse Genotyping Kit (Sigma-Aldrich, KK7302) according to the manufacturer’s instructions. PCR amplification was performed using the primers forward TTTTCACTGGATTGGGTATCAAAGG and reverse GTTGCAGTTTATTTGTGTGCCTTTTCC. All animal experiments were conducted in accordance with Swiss federal legislation on animal welfare, including the Animal Welfare Act (Loi sur la protection des animaux, LPA) and the Animal Welfare Ordinance (OPAn), and were approved under license GE15220A issued by the Cantonal Office of Health.

### 4.2 Voluntary wheel running

Experiments were conducted at the Small Animal Phenotyping Platform of the Faculty of Medicine, University of Geneva. We housed four mice per group individually in cages equipped with a running wheel, allowing voluntary physical activity. Mice had free access to the wheels throughout the experiment. We performed measurements using the TSE LabMaster system, which consists of 16 individual cages. We recorded locomotor activity at 15 min intervals over 6 consecutive days. Data are presented as the total distance travelled per mouse over the entire 6-day recording period.

### 4.3 Ex vivo muscle force measurements

Ex vivo isometric force measurements were performed on extensor digitorum longus (EDL) and soleus muscles. Animals were euthanized in accordance with institutional and national guidelines. EDL and soleus muscles were dissected bilaterally under a stereomicroscope, preserving tendon integrity at both ends, and immediately placed in continuously oxygenated (95% O₂, 5% CO₂) Krebs solution containing (in mM): 137 NaCl, 5 KCl, 1 MgSO₄, 1 NaH₂PO₄, 24 NaHCO₃, 2 CaCl₂, and 11 glucose (pH7.4) at room temperature. Muscles were mounted between a fixed hook and a force transducer (Aurora Scientific 1300A, Aurora, ON, Canada) using silk sutures tied to the tendons. Electrical field stimulation was delivered using supramaximal square-wave pulses of 0.2 ms duration. After mounting, muscles were allowed to equilibrate for 15 min, during which single twitch stimulations were applied every 30 seconds. Optimal muscle length (L₀) was determined by incrementally increasing muscle length until maximal twitch force was achieved. All subsequent recordings were performed at L₀. We obtained force-frequency relationships by stimulating muscles with 500-ms pulse trains at increasing frequencies (1, 30, 60, 90, 120, 150, 180, 210, and 240Hz). We allowed a minimum rest period of 2 minutes between consecutive stimulations to prevent fatigue. We assessed muscle fatigue by repeated tetanic stimulations at a frequency eliciting near-maximal force. EDL muscles were stimulated at 120Hz using 500-ms trains delivered every 2.5s for a total of 40 contractions, whereas soleus muscles were stimulated at 70Hz using 500-ms trains delivered every 2.5s for a total of 60 contractions. At the end of the experiments, muscles were removed from the bath, blotted dry, and weighed for normalization of force to muscle mass when indicated.

### 4.4 Muscle histology and immunofluorescence

For histology and immunofluorescence, muscles were rapidly frozen in nitrogen-cooled isopentane, and 7-µm-thick cryosections were prepared. For myosin immunostaining, sections were air-dried for 10 min at room temperature and then blocked in PBS containing 5% goat serum (Bioconcept, 2-16F00-H) and 0.3% Triton X-100. Sections were incubated sequentially with primary antibodies and corresponding fluorescent secondary antibodies, diluted in PBS supplemented with 1% BSA and 0.3% Triton X-100. Sections were mounted using Vectashield medium containing DAPI (Vector Laboratories). Images were acquired using a Zeiss Axiocam Z1 microscope. Muscle fibre type composition and cross-sectional area distribution were quantified from laminin staining using QPath software.

### 4.5 Antibodies

For fibre typing, all samples were stained with a rabbit anti-laminin antibody (Abcam, ab11575). Antibodies against myosin heavy chains IIB (mouse IgM, BF-F3), IIA (mouse IgG1, SC-71) and I (mouse IgG2b, BA-D5), developed by S. Schiaffino (Padova, Italy), and IIX (mouse IgM, 6H1), developed by C. Lucas (Sydney, Australia), were obtained from the Developmental Studies Hybridoma Bank, created by the NICHD of the NIH and maintained at the University of Iowa, Department of Biology, Iowa City, IA 52242. All antibodies were diluted 1/200. The secondary antibodies, anti-mouse IgM-Alexa 555, anti-mouse IgG1-Alexa 647, anti-mouse IgG2b-Alexa 647, and anti-rabbit-Alexa 488, are from ThermoFisher and used at 1/1000.

### 4.6 Cell Culture and Transfection

Primary mouse myoblasts were isolated as previously described by Rosenblatt et al. (1995) [22] and cultured according to the protocol reported by Cocchiararo et al. (2022) [23]. Human primary myoblasts were cultured as previously described by Brunetti et al. (2021) [24]. Briefly, human myoblasts were isolated from semitendinosus muscle biopsies obtained as surgical waste following orthopaedic procedures. All samples were collected anonymously after written informed consent and approval by the University of Geneva Ethics Committee (protocol CCER no. PB_2016-01793 (12-259)), which was accepted by the Swiss Regulatory Health Authorities and approved by the Commission Cantonale d’Éthique de la Recherche (Geneva, Switzerland). Human myoblasts were transfected with siRNA using Lipofectamine RNAiMAX (Thermo Fisher Scientific, 13778150) according to the manufacturer’s instructions, with 40 pmol of siRNA per condition. Myogenic differentiation was induced 48 h after siRNA transfection for 48 h. The siRNA targeting *STIM1L* was described by Darbellay et al. (2011) [6], and AllStars Negative Control siRNA (Qiagen) served as the control. Cells were maintained at 37°C in a humidified atmosphere containing 5% CO₂.

### 4.7 Mn^2+^ quench experiments

Human myoblasts were differentiated on 25-mm glass coverslips, whereas murine myoblasts were differentiated on plastic-bottom dishes suitable for fluorescence microscopy (Ibidi µ-Dish 35 mm, high plasticity polymer). Myotubes were loaded in the dark for 45 minutes at room temperature with 2 µM Fura-2-AM (ref. F1201, Thermo Fisher Scientific) and 0.1% Pluronic F-127 (Invitrogen, cat. no. P3000MP). After incubation, cells were washed for 10 min to allow complete dye de-esterification. Ca^2+^ signals were recorded using a Zeiss Axio Observer A1 microscope equipped with a Lambda XL illumination system (Sutter Instrument, Novato, CA, USA). Myotubes were exposed to 500 µM Mn^2+^ in a 2 mM Ca^2+^ extracellular solution containing (in mM): 135 NaCl, 5 KCl, 1 MgCl₂, 10 HEPES, 10 glucose, and 2 CaCl₂, adjusted to pH 7.4 with NaOH. Mn^2+^ was added prior to the simultaneous addition of 1 µM thapsigargin (Tg) and 10 mM caffeine, and the quenching of Fura-2 fluorescence was monitored at the isosbestic excitation wavelength of 360 nm (360BP10; Omega Optical). Images were acquired at 2 sec intervals. Emitted fluorescence was collected through a 415 DRLP dichroic mirror and a 510WB40 emission filter (Omega Optical) and detected using a cooled 16-bit sCMOS camera (pco.Edge, Visitron Systems, Puchheim, Germany). Image acquisition and analysis were performed using VisiView software (version 4.4.0.11; Visitron Systems).

### 4.8 Quantitative PCR

Total RNA was extracted from skeletal muscle using RNeasy Fibrous Tissue Mini Kit (Qiagen). DNAse-treated mRNA were reverse transcribed using High-Capacity cDNA Reverse Transcription Kit (ThermoFisher). Quantitative PCR were performed with Power Up Sybr Master Mix (ThermoFisher), analysed with StepOne software, and normalized on *Tbp* expression. qPCR amplification was performed using the primers *Stim1*-Pan-fw ACGATGCCAATGGTGATGTG and *Stim1*-Pan-rev CACCTCATCCACAGTCCAGT or *Stim1*-short-fw CCTTGTCCATGCAGTCCCC and *Stim1*-short-rev GCATGGCATTGAGAGCTTCA.

### 4.9 Western Blot

Muscle samples were snap-frozen in liquid nitrogen and mechanically pulverized into a fine powder using a hammer. Proteins were extracted by incubating the tissue lysates for 2 h at 4 °C in RIPA+ buffer containing 50 mM Tris–HCl (pH 8.0), 150mM NaCl, 1% NP-40, 0.5% sodium deoxycholate, 0.1% SDS, 1% Triton X-100, and 10% glycerol, followed by two rounds of sonication. Proteins were extracted from human myotubes using CHAPS lysis buffer (1% CHAPS, 50 mM Tris−HCl pH 7.5, 150 mM NaCl, 5 mM EDTA, 10% glycerol). Protein concentrations were determined using the Pierce BCA Protein Assay Kit (Thermo Scientific, cat. no.23225). Equal amounts of protein (15µg) were separated on precast 4-20% Bis-Tris gradient gels (mPAGE, Merck, MP42G12) and subsequently processed for immunoblotting. Proteins were transferred onto nitrocellulose membranes (Immobilon-NC, Millipore HATF85R), which were briefly blocked for 5min in polyvinyl alcohol (PVA). After washing, membranes were then incubated overnight at 4°C with primary antibodies diluted in 5% bovine serum albumin (BSA) in Tris buffered saline containing 0.1% Tween 20 (TTBS). After washing, membranes were incubated with HRP conjugated secondary antibodies diluted in 5% non fat dry milk in TTBS. Immunoreactive bands were visualised using standard chemiluminescence detection (SuperSignal West PicoPLUS, Thermoscientific 34580). Band intensities were quantified using ImageJ software, and protein expression levels were normalized to α tubulin, used as a loading control. The following primary antibodies were used: STIM1 (Millipore, cat. no. AB9870, lot3015362, rabbit polyclonal, 1:2000 dilution) and α tubulin (Sigma Aldrich, mouse monoclonal DM1A, 1:10000 dilution). Secondary antibodies included HRP-conjugated goat anti-mouse IgG (H+L) (Bio-Rad, cat. no.170 6516, 1:6000 dilution) and HRP-conjugated goat anti-rabbit IgG (H+L) (Cell Signalling Technology, lot27, 1:6000 dilution).

### 4.10 T-system Ca^2+^ handling measurements in skinned EDL fibres

T-system Ca^2+^ handling measurements were performed in EDL muscle from male WT and Stim1l-KO mice as previously described in Cully et al, 2016 [7]. Briefly, after mice euthanasia via cervical dislocation, the whole EDL muscle was rapidly dissected and placed under paraffin oil in a glass Petri dish covered with Sylgard. A bundle of fibres was mechanically dissected and exposed to a standard Ringer solution containing 2.5 mM of the low-affinity Ca^2+^-sensitive dye Rhod-5N (Invitrogen, R14207) for 10-15 min. Individual fibres were isolated from the Rhod-5N-loaded bundle and skinned by removing the sarcolemma. Skinned fibres were then transferred to a custom-made chamber filled with an intracellular solution formulated as follows: (in mM) 90 HEPES, 10 EGTA, 9.86 MgO2, 9.7 CaCO3, 8 Na2ATP, and 10 NaCP, with pH adjusted to 7.1 with KOH, resulting in a free [Ca^2+^] and free [Mg^2+^] of 67 nM and 1 mM, respectively. 4-methyl-N-(phenylmethyl) benzenesulfonamide («BTS», #1870, Tocris Bio-Techne) was dissolved in dimethyl sulfoxide (DMSO) and added at a final concentration of 50 µM to inhibit fibre contraction during the imaging protocol. Ca^2+^ depletion from the t-system was induced in a nominally Ca^2+^ and Mg^2+^-free solution with the addition of 30 mM caffeine (cat.no. C0750, Sigma). The chamber was transferred onto the stage of a Nikon Eclipse Ti2 confocal microscope for continuous Rhod-5N fluorescence recording (Nikon NIS Acquisition Software). R5N fluorescence was imaged as previously described (Koenig et al., 2019), using a Nikon 20x Plan Apo λ NA 0.75 objective on a Nikon laser scanning confocal microscope system (Nikon A1R+). Rhod-5N was excited at 561 nm at a laser power of 1%, and emitted light was collected at 595/50 nm. Minimal (Fmin) and maximal (Fmax) dye fluorescence were determined in the presence of maximal (5 mM) and nominally Ca^2+^-free solutions containing 25 µM of ionomycin (Sigma, I9657) and A23187 (Sigma, C7522). The spatially averaged real-time fluorescent signal (F) was converted to calcium concentration using the formula: [Ca^2+^] = KD × (F − Fmin)/(Fmax − F)with a KD of 0.872 mM [7].

### 4.11 Intracellular Ca^2+^ handling measurements in intact FDB fibres

Flexor digitorum brevis (FDB) fibres were enzymatically isolated following an adaptation of the protocol described in Miranda et al., 2020 [25]. Briefly, both FDB muscles were rapidly dissected from male WT and Stim1l-KO mice previously euthanized via cervical dislocation. Isolated FDB muscles were digested at 37°C for 1 hour in a MEM-based buffer supplemented with 10% FBS, 2mM L-Glutamine, 1x penicillin-streptomycin and 1000U/ml of Collagenase type II with gentle shaking. After digestion, FDB muscles were transferred to a MEM-based buffer composed as above but without Collagenase and triturated using glass pipettes of increasingly-smaller bore size in order to dissociate single fibres from the digested tissue. During this process, the fibres were monitored under a brightfield microscope, and the trituration was stopped either after 20-25 pipetting steps/bore size or when fibres began to appear fragmented. Isolated FDB fibres were then seeded in 3.5mm glass bottom petri dishes in MEM supplemented with 10% FBS, 2mM L-Glutamine, 1x penicillin-streptomycin and kept in a cell culture incubator until use. For intracellular Ca^2+^ measurements, FDB fibres were preloaded with 5 µM Fluo-4 AM (Thermo Fisher Scientific, F14201), a cell-permeable Ca^2+^-sensitive fluorescent dye, for 15 minutes at 37°C and transferred into a standard Ringer solution containing 50µM BTS to inhibit fibres contraction during the experiment. Intracellular Ca^2+^ transients were elicited via electrical field stimulation (EFS) at 1 Hz using a pair of platinum electrodes connected to a S48 GRASS stimulator, with rectangular pulses at 60V and with a duration of 4ms. Fluorescent signals (excitation at 495 nm and emission at 528 nm) were recorded using a Nikon A1R+ confocal microscope system. Perfusion of fibres with an external solution was performed using an OctaFlow II perfusion system (ALA Scientific Instruments). Minimal (Fmin) and maximal (Fmax) dye fluorescence were determined in the presence of maximal (5 mM) and nominally Ca^2+^-free Ringer buffers containing 25 µM of ionomycin (Sigma, I9657) and A23187 (Sigma, C7522). The spatially averaged real-time fluorescent signal (F) was converted to calcium concentration using the formula: [Ca^2+^] = KD × (F − Fmin)/(Fmax − F) with a KD of 1 µM [26].

## Authors’ contributions

O.D. and G.L. performed the experiments and analysed the data. E.L. performed the experiments, analysed the data, and was a major contributor to writing the manuscript. P.C., X.K. and M.F. designed and coordinated the study and wrote the manuscript. S.K. performed the experiments, analysed the data, designed and coordinated the study, and wrote the manuscript.

All authors read and approved the final manuscript.

---

**Figure S1.**
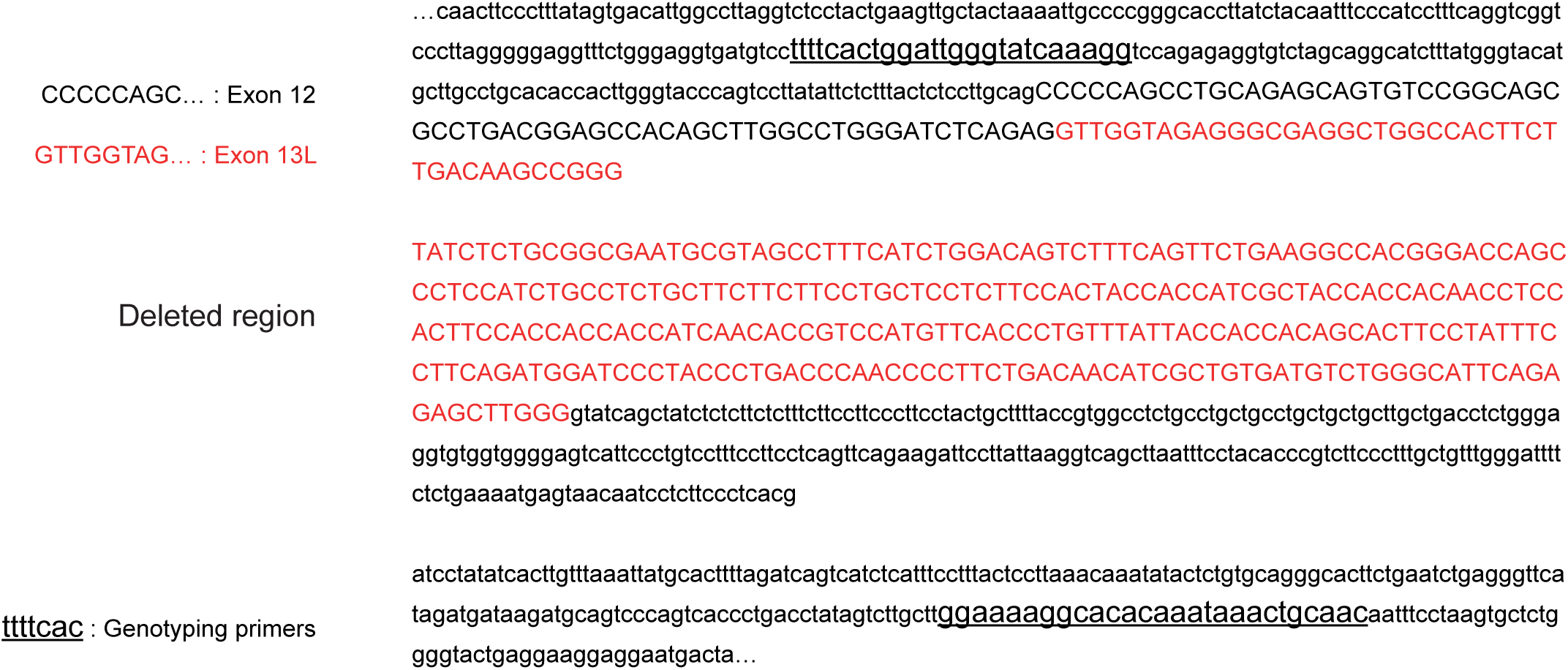
Sequence of the targeted *Stim1* locus in STIM1L KO mice. Genomic sequence of mouse *Stim1* spanning exon 12, exon 13L and the flanking introns. Exons are indicated in uppercase, while introns are shown in lowercase. Intron sequences are only partially transcribed. Exon 12 is shown in black and exon 13L in red. The 521-bp region deleted in KO mice is shown as a separate block (label “Deleted region”). It includes the 3′ 283 bp of exon 13L and the first 238 bp of the downstream intron, which removes the exon 13L splice donor site. The genotyping primers are underlined: forward 5′-TTTTCACTGGATTGGGTATCAAAGG-3′ and reverse 5′-GTTGCAGTTTATTTGTGTGCCTTTTCC-3′. They give products of [932] bp (WT allele) and [411] bp (KO allele) (see Figure 1B).

**Figure S2.**
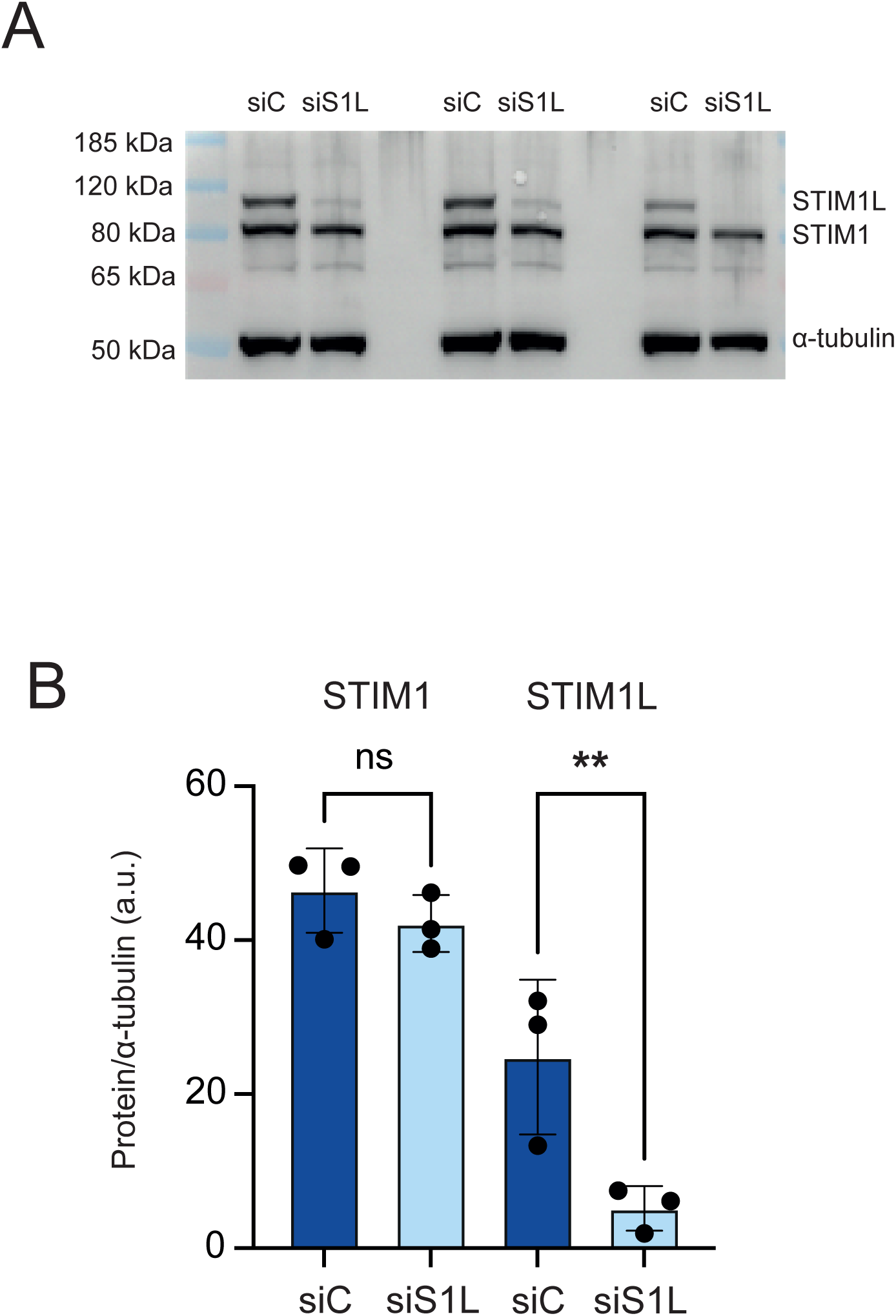
siRNA knockdown of STIM1L in human myotubes. (A) Immunoblot showing STIM1L and STIM1 in human primary myotubes transfected with control siRNA (siC) or STIM1L-specific siRNA (siS1L), using α-tubulin as the loading control. Each pair of lanes represents an independent experiment from a different donor. Molecular-weight markers are indicated on the left. (B) Quantification of STIM1 and STIM1L protein levels, normalized to α-tubulin. siSTIM1L reduced STIM1L by 79% and did not significantly change STIM1. Data are mean ± SD with individual data points. Two-way ANOVA (siRNA × protein. \*\**P* < .01; ns, not significant. siCtrl, control siRNA; siSTIM1L, STIM1L-targeting siRNA.

